# Changes in Plasma Fatty Acid Abundance Related to Chronic Pancreatitis: A Pilot Study

**DOI:** 10.1101/2023.01.05.522899

**Authors:** Kristyn Gumpper-Fedus, Olivia Crowe, Phil A. Hart, Valentina Pita-Grisanti, Ericka Velez-Bonet, Martha A. Belury, Mitchell Ramsey, Rachel M Cole, Niharika Badi, Stacey Culp, Alice Hinton, Luis Lara, Somashekar G. Krishna, Darwin L. Conwell, Zobeida Cruz-Monserrate

## Abstract

**Objectives:** Chronic pancreatitis (CP) is an inflammatory disease that affects the absorption of nutrients like fats. Molecular signaling in pancreatic cells can be influenced by fatty acids (FAs) and changes in FA abundance could impact CP-associated complications. Here, we investigated FA abundance in CP compared to controls and explored how CP-associated complications and risk factors affect FA abundance.

**Methods:** Blood and clinical parameters were collected from subjects with (n=47) and without CP (n=22). Plasma was analyzed for relative FA abundance using gas chromatography and compared between controls and CP. Changes in FA abundance due to clinical parameters were also assessed in both groups.

**Results:** Decreased relative abundance of polyunsaturated fatty acids (PUFAs) and increased monounsaturated fatty acids (MUFAs) were observed in subjects with CP in a sex-dependent manner. The relative abundance of linoleic acid increased, and oleic acid decreased in CP subjects with exocrine pancreatic dysfunction and a history of substance abuse.

**Conclusions:** Plasma FAs like linoleic acid are dysregulated in CP in a sex-dependent manner. Additionally, risk factors and metabolic dysfunction further dysregulate FA abundance in CP. These results enhance our understanding of CP and highlight potential novel targets and metabolism-related pathways for treating CP.

## Introduction

Pancreatitis is a debilitating inflammatory disease of the pancreas, which can occur as an acute or chronic disease. Risk factors for chronic pancreatitis (CP) include excessive alcohol consumption, smoking, genetic predisposition, autoimmune disease, and ductal obstructions.^1^ CP is characterized by inflammation of increasing severity, resulting in declining pancreatic function and irreversible morphologic changes, including calcification and fibrosis.^2^ The unremitting abdominal pain that accompanies this fibro-inflammatory process is linked to reduced quality of life, increased disability, and increased utilization of healthcare services.^3^ In addition, CP is a recognized risk factor for pancreatic ductal adenocarcinoma (PDAC), a highly aggressive malignancy that is difficult to detect at an early stage.^2,4,5^

Nutritional complications are some of the hallmarks of CP due to diabetes and exocrine pancreatic dysfunction (EPD), including impaired fat absorption secondary to pancreatic lipase deficiency. These sequelae lead to direct and indirect consequences, including altered dietary patterns, that lead to changes in fatty acid (FA) metabolism. In prior studies from Europe and Japan, patients with CP exhibited differences in FA abundance compared with healthy individuals.^6,7^ Moreover, others found that diabetes, when combined with CP, further exacerbates alterations in the abundance of plasma FAs.^8^ Additionally, changes in FA abundance have been associated with increased adiposity, a driver of diabetes, which is highly prevalent in CP.^9-11^ FA supplementation with omega-3 polyunsaturated fatty acids (PUFAs), such as eicosapentaenoic acid and docosapentaenoic acid, has been explored as a therapeutic strategy to reduce the severity of inflammation in pancreatitis; however, results are mixed.^12,13^ Additionally, these studies do not address whether sex or age also influence FA abundance in CP.

In this pilot study, we aimed to compare changes in FA abundance between CP and healthy individuals. We further explored associations between FA abundance and sex, age, metabolic complications, including changes in body mass index (BMI), diabetes, and EPD, and known risk factors for CP (alcohol consumption and smoking) in subjects with CP.

## Methods

### Study Population

Healthy control subjects (n=22) and subjects with CP (n=47) were enrolled in an observational study conducted at The Ohio State University Wexner Medical Center Pancreas Clinic from 2015 to 2016, which was approved by the OSU Institutional Review Board. Ten patients with CP did not have pancreatic calcifications but met other diagnostic criteria for CP.^2^ Control subjects included individuals with a family history of PDAC but did not have imaging or clinical evidence of CP or other diseases of the exocrine pancreas.

### Clinical Data

Clinical data were obtained by reviewing medical records and recorded on a standardized form. Extracted information included disease history, past medical history, and social history. Smoking status and alcohol use patterns were self-reported prospectively using a standardized questionnaire. Excessive alcohol use was defined as consuming ≥14 alcoholic drinks per week for ≥5 years and tobacco use as smoking ≥100 cigarettes during the subject’s lifetime. EPD was defined as the presence of a supporting clinical diagnosis (based on either overt steatorrhea or abnormal indirect pancreatic function testing) and/or the use of pancreatic enzyme replacement therapy.

### Blood Collection and Processing

Non-fasting blood samples were collected from all study subjects at the time of enrollment in the study. Aliquots of plasma were frozen at -80°C until the time of sample analysis. Research personnel conducting the FA analyses were blinded to the subjects’ group assignment.

### Fatty Acid Analysis

Lipids were extracted from formerly frozen plasma samples using the 2:1 chloroform:methanol method, ^14,15^ as we previously described.^16^ Briefly, samples were washed with 0.88% potassium chloride, followed by FA methyl ester extraction with hexane after being prepared with 5% hydrochloric acid in methanol at 76°C. FA methyl esters were analyzed using gas chromatography on a 30-m Omegawax TM 320 fused silica capillary column (Supelco Inc, Bellefonte, PA, USA), with the oven temperature and carrier gas flow rate settings as previously described.^17^ Samples were compared to standards for each FA methyl ester (Supelco Inc; Matreya, LLC, Pleasant Gap, PA; Nu-Check Prep Inc, Elysian, MN). The data were presented as the quantity of each FA as a percentage of all identified FAs (% area) indicating the relative abundance of each fatty acid, the most common way of analyzing fatty acid composition.^17,18^ Using this method, docosahexaenoic acid and nervonic acid (24:1n9) co-eluted during chromatography. Because nervonic acid typically represents < 1% of plasma FAs, all data reported for docosahexaenoic acid also includes nervonic acid. The percent coefficient of variance (%CV) for the runs is provided for inter-sample variability calculation (**Supplement Table 1**). We also grouped FAs by saturation type to compare relative abundance among CP vs controls.

**Table 1.** Characteristics of control and CP groups.

|  | **Control**<br>**(n = 22)** | **CP**<br>**(n = 47)** | **p-value** |
| --- | --- | --- | --- |
| Age in years, mean (SD) | 46.6 (15.2) | 54.8 (14.7) | **0.036** |
| Male sex, n (%) | 7 (31.8) | 31 (66.0) | **0.010** |
| BMI, mean (SD) | 29.5 (6.6) | 23.9 (4.8) | **<0.001** |
| Diabetes present, n (%) | 1 (4.6) | 21 (55.3) | **0.001** |
| EPD present, n (%) | 0 (0.0) | 29 (61.7) | **<0.001** |
| Alcohol Use ($\geq 14$ drinks/week), n (%) | 3 (14.3)* | 21 (48.8)* | **0.012** |
| Smoking, n (%) | 10 (45.5) | 30 (63.8) | 0.194 |

\* Data not available for all individuals, SD = Standard Deviation

### Statistical Analysis

Statistical analyses were performed using GraphPad Prism 9 (GraphPad Software Inc, San Diego, CA). Frequencies were reported for categorical variables and were compared between groups using a chi-squared test of independence or Fisher’s exact test, as appropriate. Means and standard deviations were reported by study group for the continuous variables of age and BMI and compared using an independent samples t-test and the non-parametric Mann-Whitney U test, respectively. Differences in FA relative abundance between groups were compared using 2-way ANOVA with a Fisher’s LSD post-hoc test for pairwise comparisons. A non-parametric Mann-Whitney U test was used for FAs whose distribution was non-normal. P-values were adjusted for multiple testing using a Bonferroni correction. Spearman’s rho correlation coefficients were calculated to assess associations between FA relative abundance and age or BMI.

## Results

### Study Population

A total of 69 subjects were included in the study population: 22 controls and 47 subjects with CP. Subjects with CP were more likely to be male, older, and have a lower BMI (**Table 1**). The CP group had a higher prevalence of diabetes and excessive alcohol use compared with the control group, but no difference was detected for history of smoking.

### Relative Abundance of FAs Are Altered in CP in a sex-dependent manner

When classifying FAs by carbon chain saturation type, we found an increase in monounsaturated FAs (MUFAs) and a decrease in PUFAs the CP group (**Figure 1A, Supplemental Table 1**). There was no difference in overall relative abundance of saturated fatty acids (SFAs). Since there was a significant difference in sex between the control and CP groups, we compared relative abundances of FA between control and CP groups in males and females separately. There was no overall significant change in carbon chain saturation types in the males (**Figure 1B**), however the MUFA relative abundance was elevated and PUFA relative abundance was decreased in the females with CP compared to controls (**Figure 1C)** suggesting that the effects observed in (**Figure 1A**) could be driven by the females.

**Figure 1.**
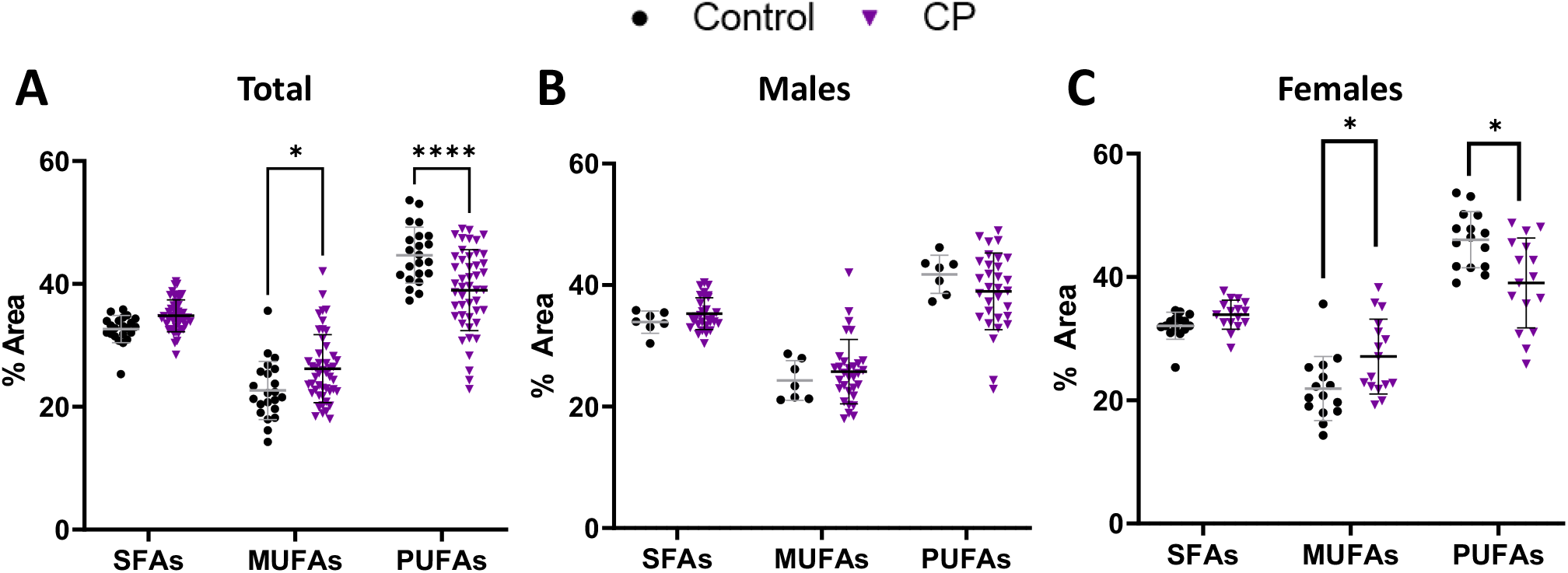
FA relative abundance grouped by carbon chain saturation type between subjects with CP and controls. Relative abundance (% area) of FAs grouped by carbon chain saturation type were compared between (**A**) all controls (n=22) and subjects with CP (n=47), (**B**) male controls (n=7) and males with CP (n=31), and (**C**) female controls (n=15) and females with CP (n=16). Significance was determined using t-tests with a Bonferroni correction for 3 comparisons. *p<0.05. **p<0.01 and ****p<0.0001. CP, chronic pancreatitis; SFAs, saturated fatty acids; MUFAs, monounsaturated fatty acids; PUFAs, polyunsaturated fatty acids

When the relative abundance of each FAs was analyzed, the abundance of the PUFA linoleic acid was significantly lower and the abundance of the SFA palmitic acid was significantly elevated in CP compared to controls (**Figure 2A**). In males, only a decrease in linoleic acid was observed, however the abundance of palmitic acid trended to be higher (unadjusted p=0.0445) (**Figure 2B, Supplemental Table 3**). In addition to a significant decrease in linoleic acid in females with CP, the relative abundance of palmitic acid was also elevated in females with CP compared to controls (**Figure 2C**). There was a trend of increased palmitoleic (unadjusted p=0.0055), oleic (unadjusted p=0.0192), and vaccenic acids (unadjusted p=0.0459) in females with CP compared to controls, suggesting that the changes in MUFAs in females with CP might drive the overall relative abundance changes when separating by carbon chain saturation type (**Supplemental Table 4**).

**Figure 2.**
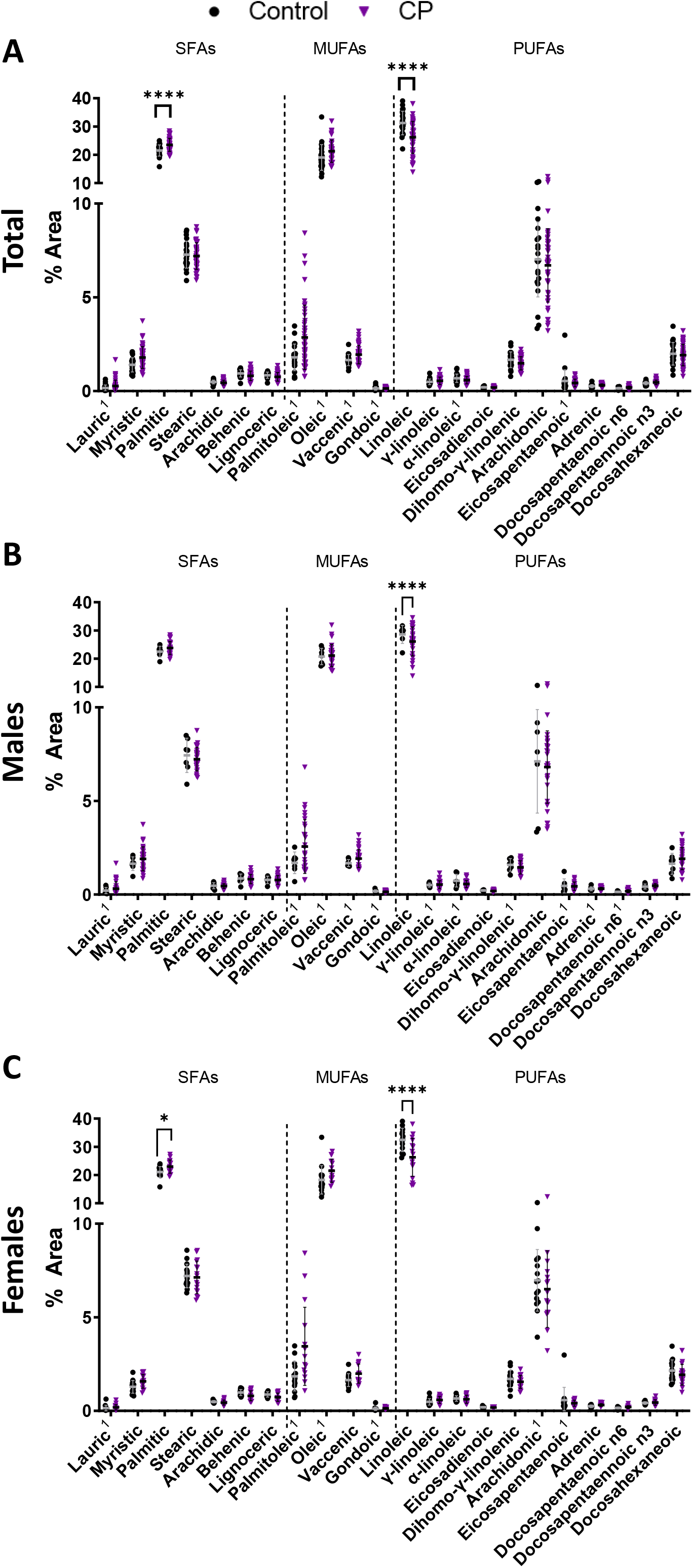
Relative abundance of individual FAs between subjects with CP and controls. Relative abundance (% area) of FAs (**A**) all controls and subjects with CP, (**B**) males (control n=7, CP n=31), and (**C**) females (control n=15, CP n=16). Significance was determined using t-tests with a Bonferroni correction for 22 comparisons. **p<0.01 and ****p<0.0001. ^1^Non-parametric Mann-Whitney test used for non-normal data. CP, chronic pancreatitis; SFAs, saturated fatty acids; MUFAs, monounsaturated fatty acids; PUFAs, polyunsaturated fatty acids

### Most FAs Are Not Correlated with Age and BMI in CP

Since age and BMI characteristics were different between the control and CP groups and can influence FA metabolism,^19,20^ we explored the changes in FAs composition based on these characteristics. Lauric, eicosadienoic, and dihomo-γ-linolenic acids were negatively correlated with age while oleic and docosapentaenoic were positively correlated with age in control subjects (**Table 2**). These correlations were not detected in subjects with CP. In controls, SFAs were positively correlated with BMI while PUFAs trended toward a negative correlation with BMI (**Table 3**). Additionally, there was a positive correlation between BMI and the relative abundance of myristic, palmitic, palmitoleic, vaccenic, and dihomo-γ-linolenic acids in the control group (**Table 3**). Like age, correlations between BMI and most of the FAs were not detected in the CP group; however, arachidic and docosahexanoic acids remained negatively correlated with BMI in both groups.

**Table 2.** Correlations between individual FAs and Age in the control and CP groups.

| *Fatty Acid* | *Control*<br>(*n*=22) |  | *CP*<br>(*n*=47) |  |
| --- | --- | --- | --- | --- |
|  | **r** | **p-value** | **r** | **p-value** |
| ***SFAs*** | -0.0023 | 0.9920 | 0.1808 | 0.2239 |
| *Lauric* | **-0.4151** | **0.0547** | -0.0241 | 0.8723 |
| *Myristic* | 0.0915 | 0.6854 | 0.0297 | 0.8431 |
| *Palmitic* | -0.0254 | 0.9106 | 0.1557 | 0.2961 |
| *Stearic* | -0.0441 | 0.8457 | 0.1113 | 0.4564 |
| *Arachidic* | 0.1330 | 0.5553 | 0.0867 | 0.5625 |
| *Behenic* | -0.1231 | 0.5851 | -0.0122 | 0.9354 |
| *Lignoceric* | -0.1425 | 0.5269 | -0.0457 | 0.7604 |
| ***MUFAs*** | 0.3868 | 0.0754 | 0.1175 | 0.4313 |
| *Palmitoleic* | -0.0593 | 0.7932 | -0.0758 | 0.6128 |
| *Oleic* | **0.4184** | **0.0526** | 0.1477 | 0.3217 |
| *Vaccenic* | 0.1915 | 0.3933 | -0.0512 | 0.7326 |
| *Gondoic* | -0.0302 | 0.8938 | 0.1420 | 0.3410 |
| ***PUFAs*** | -0.3348 | 0.1277 | -0.1133 | 0.4484 |
| *Linoleic* | **-0.5776** | **0.0049** | -0.1163 | 0.4362 |
| *γ-linoleic* | 0.1668 | 0.4582 | 0.0572 | 0.7025 |
| *α-linoleic* | 0.0814 | 0.7189 | -0.0857 | 0.5666 |
| *Eicosadienoic* | **-0.4387** | **0.0411** | -0.0840 | 0.5748 |
| *Dihomo-γ-linolenic* | **-0.4270** | **0.0475** | 0.0917 | 0.5397 |
| *Arachidonic* | 0.3032 | 0.1701 | 0.0514 | 0.7318 |
| *Eicosapentaenoic* | 0.3616 | 0.0982 | 0.1532 | 0.3038 |
| *Adrenic* | 0.1500 | 0.5053 | -0.0876 | 0.5584 |
| *Docosapentaenoic n6* | -0.0228 | 0.9198 | -0.0970 | 0.5165 |
| *Docosapentaenoic n3* | **0.4908** | **0.0204** | -0.0658 | 0.6605 |
| *Docosahexaneic* | 0.1452 | 0.5191 | 0.0457 | 0.7604 |

Bolded values are p<0.05. CP, chronic pancreatitis.

**Table 3.** Correlations between individual FAs and BMI in the control and CP groups.

| Fatty Acid | Control<br>(n=22) |  | CP<br>(n=47) |  |
| --- | --- | --- | --- | --- |
|  | r | p-value | r | p-value |
| **SFAs** | **0.5874** | **0.0040** | -0.0426 | 0.7764 |
| Lauric | 0.1880 | 0.4021 | -0.1587 | 0.2867 |
| Myristic | 0.5034 | **0.0169** | -0.0182 | 0.9033 |
| Palmitic | **0.7143** | **0.0002** | 0.1310 | 0.3800 |
| Stearic | 0.0525 | 0.8164 | -0.1879 | 0.2060 |
| Arachidic | -0.4045 | 0.0618 | -0.2468 | 0.0945 |
| Behenic | -0.3423 | 0.1189 | -0.2068 | 0.1630 |
| Lignoceric | **-0.4960** | **0.0189** | -0.2468 | 0.0945 |
| **MUFAs** | 0.1835 | 0.4137 | 0.1646 | 0.2689 |
| Palmitoleic | **0.5637** | **0.0063** | -0.0578 | 0.6998 |
| Oleic | 0.1756 | 0.4344 | 0.2331 | 0.1149 |
| Vaccenic | 0.3886 | 0.0739 | -0.0621 | 0.6786 |
| Gondoic | 0.0525 | 0.8167 | 0.1268 | 0.3959 |
| **PUFAs** | -0.3868 | 0.0754 | -0.0993 | 0.5067 |
| Linoleic | -0.1925 | 0.3906 | -0.0722 | 0.6298 |
| γ-linoleic | 0.2340 | 0.2945 | 0.0719 | 0.6310 |
| α-linoleic | 0.1740 | 0.4386 | -0.0323 | 0.8292 |
| Eicosadienoic | 0.1955 | 0.3832 | -0.1105 | 0.4598 |
| Dihomo-γ-linolenic | **0.7100** | **0.0002** | -0.0764 | 0.6097 |
| Arachidonic | -0.3495 | 0.1108 | 0.0265 | 0.8598 |
| Eicosapentaenoic | -0.2305 | 0.3020 | 0.0023 | 0.9877 |
| Adrenic | 0.3237 | 0.1417 | 0.0760 | 0.6115 |
| Docosapentaenoic n6 | 0.1317 | 0.5592 | -0.1065 | 0.4761 |
| Docosapentaenoic n3 | 0.0493 | 0.8277 | 0.1573 | 0.2911 |
| Docosahexaneic | **-0.4960** | **0.0150** | **-0.2833** | **0.0536** |

Bolded values are p<0.05. CP, chronic pancreatitis.

### Comorbidities and Risk Factors for CP Influence the Relative Abundance of FAs

Diabetes and EPD are common comorbidities in CP that influence nutrition absorption and metabolism.^21^ First, subjects with CP were categorized according to the presence or absence of diabetes (**Table 1**). There was no between group differences in FA carbon chain saturation types when subjects with CP were dichotomized by the presence or absence of diabetes (**Supplemental Table 5**). However, the relative abundance of oleic acid was decreased, and linoleic acid was increased in subjects with CP and diabetes compared to subjects with CP without diabetes (**Figure 3A and Supplemental Table 5**). Additionally, there was a trend increase in lauric acid in subjects with CP and diabetes compared to those without diabetes (unadjusted p=0.0417) (**Figure 3A and Supplemental Table 5**).

**Figure 3.**
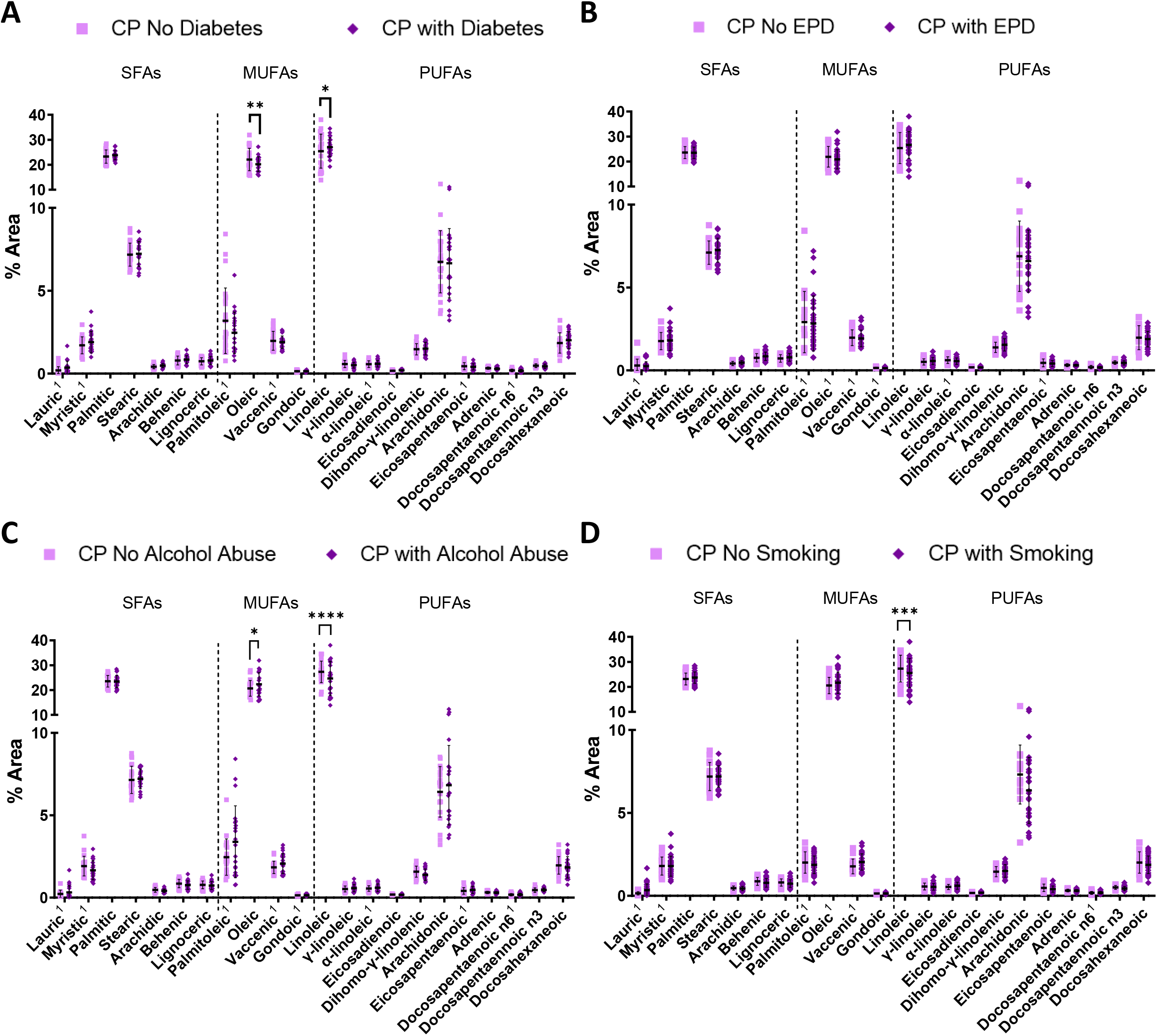
Relative abundance of FAs associated with comorbidities and risk factors for CP. Relative abundance (% area) of individual FAs comparing CP subjects by (**A**) with diabetes (n=21) and without diabetes (n=26), (**B**) with EPD (n=29) and without EPD (n=18), (**C**) alcohol abuse (n=21) or no alcohol abuse (n=22), and (**D**) smoking (n=30) or no smoking (n=17). Significance was determined using two-way ANOVA, followed by Bonferroni correction for 22 comparisons. *p<0.05, **p<0.01, ***p<0.001, p<0.0001. ^1^Non-parametric Mann-Whitney test used for non-normal data. CP, chronic pancreatitis; SFAs, saturated fatty acids; MUFAs, monounsaturated fatty acids; PUFAs, polyunsaturated fatty acids

EPD alters nutrient absorption, so we looked at whether EPD affected plasma FA relative abundance in CP. There was no difference in the relative abundance of FA carbon chain saturation types when subjects with CP were dichotomized by the presence or absence of EPD (**Supplemental Table 6**). Additionally, there was no difference in the relative abundance of individual FAs when subjects with CP were dichotomized by the presence or absence of EPD; however, linoleic acid exhibited an increased trend in subjects with CP and EPD (unadjusted p=0.0143) (**Figure 3B, Supplemental Table 6**). This suggests that only linoleic acid maybe affected by EPD in CP.

Excessive tobacco and alcohol use are common risk factors for CP.^22^ Therefore, we evaluated whether changes in the relative abundance of FAs was associated with substance use and CP. We found no difference in the relative abundance of FAs when comparing carbon chain saturation types in CP based on a history of alcohol use (**Supplemental Table 7**). Linoleic acid relative abundance was decreased, and oleic acid relative abundance was increased in subjects with CP who had a history of alcohol use (**Figure 3C, Supplemental Table 7**). We also found no difference in FA relative abundance when comparing carbon chain saturation types in subjects with CP based on of a history of smoking (**Supplemental Table 8**); however linoleic acid was also decreased in subjects with CP who had a history of smoking (**Figure 3D, Supplemental Table 8**). There was also a trend for a decreased relative abundance of arachidonic acid in subjects with CP who had a history of smoking (unadjusted p=0.0558) (**Supplemental Table 8**).

## Discussion

Here, we show that plasma FA methyl ester relative abundance is different between CP and healthy controls and there was a different magnitude of effects based on sex. Additionally, correlations normally observed between FAs and BMI or age are lost in CP. We also demonstrated that plasma FA relative abundance (particularly linoleic acid) is altered by pancreatic insufficiency (diabetes or EPD) or a history of substance use (alcohol and smoking) in CP. Overall, these results provide evidence for further investigation into the mechanisms that drive FA changes in CP and whether this may identify potential therapeutic targets.

Chronic inflammation, as observed in CP, leads to increased production of reactive oxygen or nitrogen species, which react with membrane lipids, such as linoleic acid, and result in an increased relative abundance of oxidated FAs that can lead to cell damage and death.^23^ Additionally, palmitic acid, which was upregulated in the female CP group, can induce inflammation by stimulating secretion of pro-inflammatory cytokines like interleukin 6, tumor necrosis factor α, interleukin 8, and interleukin 1β, inducing insulin resistance.^24,25^ Diets high in palmitic and myristic acids have been associated with lipotoxicity related to non-alcoholic steatohepatitis.^26^ Although we observed a similar decrease in linoleic acid to other studies,^6^ we also observed sex-specific changes in the relative abundance of MUFAs and PUFAs in female CP patients; however only a trend in MUFA and PUFA abundance in males. An assessment of the urine proteome after total pancreatectomy with islet autotransplantation also indicated a strong difference between males and females.^27^ Moreover, a study using a large cohort from Australia that assess risk factors for diabetes and cardiovascular disease, identified sex-specific differences in lipid classes and species.^28^ Our data suggests that sex may also influence some of the lipid-associated metabolic dysfunction seen in pancreatic diseases like CP. However, this will need to be validated in future studies using a larger sample size and from multiple centers since the sample size of control males in the study population was lower. Additionally, PUFAs like the omega-3 eicosapentaenoic and docosahexanoic acids are often recommended as nutritional supplementation in the form of fish oil to reduce inflammation in acute pancreatitis.^12,13^ However, we did not observe dysregulation of these specific PUFAs in CP. Further assessment of the potential antioxidant benefit of these supplements or whether targeting linoleic acid supplementation or metabolism could improve CP-related inflammation is warranted.

Increased age generally correlates with changes in FA composition.^29-31^ Additionally, older patients and those with CP often have increased visceral adipose tissue and peripancreatic fat, compared with individuals without CP, even though CP patients tend to be leaner.^32,33^ Fatty replacement of pancreatic acinar cells and inflammation from increased adipokine levels can lead to exocrine dysfunction.^34^ Although visceral adiposity is increased in CP and aging,^32,33^ the correlations observed between age or BMI and the relative abundance of several FAs in the control group did not persist in the CP group. To our knowledge, this is the first study to identify a loss of correlation between plasma FA abundance and age and BMI in CP, but a larger sample size is needed to verify this observation. Nevertheless, this illustrates how pancreatic diseases can dysregulate metabolism and nutrient absorption beyond changes cause by differences in body composition or age.

Diabetes is a common co-morbidity of CP that contributes to metabolic dysfunction.^3,35,36^ Several studies have observed changes in FA abundance due to diabetes, including an increase in linoleic acid.^37^ However, we observed a decrease in oleic acid abundance and an increase in linoleic acid abundance in CP with diabetes compared to CP without diabetes. Oleic acid supplementation has been shown to improve insulin sensitivity of adipocytes.^38^ Additionally, high linoleic acid abundance is also associated with improved glycemic control and a reduced incidence of diabetes and reduces diabetes in preclinical models.^10^ Linoleic acid supplementation increases the expression of beneficial tetralinoleoyl cardiolipins (important in mitochondrial function) in peripheral blood mononuclear cells (PBMCs) of healthy subjects. Our finding of higher linoleic acid and lower oleic acid abundance in CP with diabetes suggests the combined endocrine and exocrine dysfunction may be modifying these FAs in a previously unreported fashion within CP. Therefore, further assessment of FAs and their metabolism in CP and diabetes together will be needed to understand the complex regulation of FAs in these disorders combined.

Substance abuse is another strong etiologic risk factor for developing CP. Consumption of more than 4–5 alcoholic drinks per day is a common etiologic factor, accounting for up half of all CP cases.^2,39^ Smoking is another independent risk factor for CP with a dose-dependent relationship.^2,22,40^ Several studies have shown FAs like linoleic acid as well as other fatty acids in its metabolic pathway are dysregulated in patients with CP who had a history of excessive alcohol consumption compared to healthy controls.^7,41,42^ In our study, the relative abundance of linoleic acid was decreased and oleic acid was increased due to alcohol use, and only linoleic acid was affected by a history of smoking in CP. Linoleic acid metabolism can result in an increase of prostaglandins that promote inflammation.^43,44^ A decrease in linoleic acid may indicate increased conversion of linoleic acid to pro-inflammatory prostaglandins, driving the pathogenesis of CP; however, future studies will need to explore this relationship in more detail.

There are limitations to this pilot study that should be considered. First, our findings, particularly those related to subgroup analyses (even with the proper statistical comparison correction) should be considered preliminary because of the small sample size, limiting the power of our analysis, and likely contributing to the overall differences observed between the control and CP groups. For example, we were unable to compare relative FA abundance in controls with diabetes or a history of substance use. Additionally, blood samples were obtained while patients were non-fasting, and nutritional intake was not recorded, which may have contributed to variances in our data compared to others did.^7,8,41^ We were also unable to assess FA changes due to other potential metabolic disorders such as hyperlipidemia.

Despite the stated limitations, the results of this pilot study show potential sex-dependent perturbations in FA relative abundance in CP. Our results highlight the intrinsic heterogeneity within CP, and reveals linoleic acid to be consistently dysregulated in subgroups within CP. Therefore, this pilot study provides the basis for larger studies using carefully processed and clinically characterized biological samples to evaluate the role of FAs and FA metabolism in the pathogenesis of CP.

