## Supplemental Tables for "Changes in Plasma Fatty Acid Abundance Related to Chronic Pancreatitis: A Pilot Study"

| Fatty acid | Omega Nomenclature | Saturation Type | %CV |
|---|---|---|---|
| Lauric acid | C12:0 | SFA | 19.39 |
| Myristic acid | C14:0 | SFA | 9.78 |
| Palmitic acid | C16:0 | SFA | 0.52 |
| Palmitoleic acid | C16:1n7 | MUFA | 2.48 |
| Stearic acid | C18:0 | SFA | 1.85 |
| Oleic acid | C18:1n9 | MUFA | 1.18 |
| Vaccenic acid | C18:1n7 | MFA | 1.26 |
| Linoleic acid | C18:2n6 | PUFA | 0.89 |
| γ-linoleic acid | C18:3n6 | PUFA | 2.13 |
| α-linolenic acid | C18:3n3 | PUFA | 1.71 |
| Arachidic acid | C20:0 | SFA | 12.64 |
| Gondoic acid | C20:1n9 | MUFA | 11.35 |
| Eicosadienoic acid | C20:2n6 | PUFA | 15.20 |
| Dihomo-γ-linolenic acid | C20:3n6 | PUFA | 1.12 |
| Arachidonic acid | C20:4n6 | PUFA | 0.55 |
| Eicosapentaenoic acid | C20:5n3 | PUFA | 7.30 |
| Behenic acid | C22:0 | SFA | 7.24 |
| Adrenic acid | C22:4n6 | PUFA | 1.93 |
| Docosapentaenoic acid n6 | C22:5n6 | PUFA | 11.69 |
| Docosapentaenoic acid n3 | C22:5n3 | PUFA | 2.89 |
| Lignoceric acid | C24:0 | SFA | 7.39 |
| Docosahexaenoic acid | C24:6n3 | PUFA | 1.91 |

**Supplemental Table 1.** Percent coefficient of variance (%CV) for each fatty acid measured

**Supplemental Table 2.** Comparison of plasma fatty acids abundance (% Area) between control and CP subjects

| Fatty Acids | Control (n=22) Mean (St. Dev) | CP (n=47) Mean (St. Dev) | Unadjusted p-value | Adjusted p-value |
|---|---|---|---|---|
| ***SFAs***[1] | 32.66 (2.22) | 34.81 (2.60) | 0.0873 | 0.2618 |
| Lauric[1] | 0.17 (0.16) | 0.27 (0.58) | 0.1935 | 0.9912 |
| Myristic | 1.40 (0.38) | 1.79 (0.58) | 0.3475 | 0.9999 |
| Palmitic | 21.54 (1.99) | 23.52 (2.32) | **<0.0001** | **<0.0001** |
| Stearic | 7.28 (0.72) | 7.20 (0.71) | 0.8509 | 1.0000 |
| Arachidic | 0.48 (0.10) | 0.45 (0.13) | 0.9339 | 1.0000 |
| Behenic | 0.94 (0.19) | 0.82 (0.25) | 0.7625 | 1.0000 |
| Lignoceric | 0.85 (0.16) | 0.76 (0.23) | 0.8389 | 1.0000 |
| ***MUFAs***[1] | 22.66 (4.74) | 26.20 (5.55) | **0.0052** | **0.0155** |
| Palmitoleic[1] | 1.80 (0.71) | 2.86 (1.72) | **0.0140** | 0.2669 |
| Oleic[1] | 19.06 (4.54) | 21.26 (3.96) | **0.0228** | 0.3984 |
| Vaccenic[1] | 1.66 (0.30) | 1.94 (0.49) | **0.0181** | 0.3311 |
| Gondoic[1] | 0.14 (0.09) | 0.14 (0.04) | 0.1405 | 0.9642 |
| ***PUFAs***[1] | 44.68 (4.55) | 38.98 (6.58) | **<0.0001** | **<0.0001** |
| Linoleic | 31.24 (4.19) | 26.17 (5.66) | **<0.0001** | **<0.0001** |
| γ-linoleic[1] | 0.49 (0.15) | 0.55 (0.20) | 0.8916 | 1.0000 |
| α-linoleic[1] | 0.68 (0.22) | 0.58 (0.20) | **0.0541** | 0.7058 |
| Eicosadienoic | 0.20 (0.04) | 0.18 (0.04) | 0.9666 | 1.0000 |
| Dihomo-γ-linolenic[1] | 0.17 (0.42) | 1.49 (0.35) | 0.6595 | 1.0000 |
| Arachidonic | 7.02 (2.00) | 6.71 (1.96) | 0.4518 | 1.0000 |
| Eicosapentaenoic[1] | 0.54 (0.59) | 0.43 (0.20) | 0.9924 | 1.0000 |
| Adrenic[1] | 0.27 (0.09) | 0.31 (0.07) | 0.9106 | 1.0000 |
| Docosapentaenoic n6[1] | 0.15 (0.05) | 0.19 (0.08) | 0.9391 | 1.0000 |
| Docosapentaenoic n3 | 0.42 (0.09) | 0.46 (0.12) | 0.9274 | 1.0000 |
| Docosahexaneoic | 1.99 (0.61) | 1.92 (0.57) | 0.8673 | 1.0000 |

[1]Non-parametric Mann-Whitney Test.
Bolded = $p<0.05$

**Supplemental Table 3.** Comparison of plasma fatty acids abundance (% Area) between male control and CP subjects

| Fatty Acids | Control (n=7) Mean (St. Dev) | CP (n=31) Mean (St. Dev) | Unadjusted p-value | Adjusted p-value |
|---|---|---|---|---|
| ***SFAs*[1]** | 33.92 (1.84) | 35.29 (2.61) | 0.3711 | 0.7513 |
| Lauric[1] | 0.21 (0.17) | 0.31 (0.37) | 0.9633 | 1.0000 |
| Myristic | 1.65 (0.36) | 1.90 (0.64) | 0.6899 | 1.0000 |
| Palmitic | 22.51 (1.93) | 23.79 (2.29) | **0.0445** | 0.6327 |
| Stearic | 7.44 (0.92) | 7.23 (0.63) | 0.7522 | 1.0000 |
| Arachidic | 0.47 (0.16) | 0.45 (0.13) | 0.9764 | 1.0000 |
| Behenic | 0.86 (0.24) | 0.82 (0.25) | 0.9542 | 1.0000 |
| Lignoceric | 0.78 (0.20) | 0.78 0.24) | 0.9948 | 1.0000 |
| ***MUFAs*[1]** | 24.31 (3.27) | 25.74 (5.30) | 0.6852 | 0.9688 |
| Palmitoleic[1] | 1.69 (0.58) | 2.56 (1.44) | 0.2690 | 0.9990 |
| Oleic[1] | 20.75 (2.88) | 21.12 (3.94) | 0.9640 | 1.0000 |
| Vaccenic[1] | 1.68 (0.19) | 1.92 (0.48) | 0.2773 | 0.9992 |
| Gondoic[1] | 0.18 (0.07) | 0.14 (0.04) | 0.1405 | 0.9642 |
| ***PUFAs*[1]** | 41.77 (3.14) | 38.96 (6.32) | 0.3139 | 0.9770 |
| Linoleic | 28.62 (3.19) | 26.11 (5.14) | **<0.0001** | **<0.0001** |
| γ-linoleic[1] | 0.51 (0.11) | 0.53 (0.21) | 0.9633 | 1.0000 |
| α-linoleic[1] | 0.74 (0.34) | 0.56 (0.19) | 0.1757 | 0.9858 |
| Eicosadienoic | 0.22 (0.04) | 0.19 (0.04) | 0.9565 | 1.0000 |
| Dihomo-γ-linolenic[1] | 1.59 (0.33) | 1.45 (0.36) | 0.3496 | 0.9999 |
| Arachidonic | 7.12 (2.77) | 6.81 (1.93) | 0.6311 | 1.0000 |
| Eicosapentaenoic[1] | 0.46 (0.37) | 0.44 (0.22) | 0.9751 | 1.0000 |
| Adrenic[1] | 0.31 (0.11) | 0.31 (0.07) | 0.7056 | 1.0000 |
| Docosapentaenoic n6[1] | 0.14 (0.04) | 0.18 (.08) | 0.4878 | 1.0000 |
| Docosapentaenoic n3 | 0.43 (0.14) | 0.47 (0.11) | 0.3685 | 1.0000 |
| Docosahexaneoic | 1.64 (0.56) | 1.92 (0.59) | 0.6689 | 1.0000 |

[1]Non-parametric Mann-Whitney Test.
Bolded = p<0.05

**Supplemental Table 4.** Comparison of plasma fatty acids abundance (% Area) between female control and CP subjects

| Fatty Acids | Control (n=15) Mean (St. Dev) | CP (n=16) Mean (St. Dev) | Unadjusted p-value | Adjusted p-value |
|---|---|---|---|---|
| ***SFAs*[1]** | 32.07 (2.19) | 33.88 (2.37) | **0.0322** | 0.0934 |
| Lauric[1] | 0.15 (0.15) | 0.19 (0.14) | 0.1978 | 0.9922 |
| Myristic | 1.28 (0.34) | 1.58 (0.37) | 0.6335 | 1.0000 |
| Palmitic | 21.09 (1.91) | 23.00 (2.34) | **0.0020** | **0.0431** |
| Stearic | 7.20 (0.63) | 7.31 (0.87) | 0.9063 | 1.0000 |
| Arachidic | 0.49 (0.07) | 0.44 (0.13) | 0.9381 | 1.0000 |
| Behenic | 0.98 (0.16) | 0.80 (0.27) | 0.7747 | 1.0000 |
| Lignoceric | 0.88 (0.14) | 0.73 (0.22) | 0.8131 | 1.0000 |
| ***MUFAs*[1]** | 21.89 (5.21) | 27.10 (6.07) | **0.0139** | **0.0412** |
| Palmitoleic[1] | 1.85 (0.78) | 3.45 (2.09) | **0.0055** | 0.1147 |
| Oleic[1] | 18.28 (5.02) | 21.52 (4.10) | **0.0192** | 0.3468 |
| Vaccenic | 1.65 (0.34) | 1.99 (0.51) | 0.5765 | 1.0000 |
| Gondoic[1] | 0.12 (0.09) | 0.13 (0.05) | 0.0682 | 0.7885 |
| ***PUFAs*[1]** | 46.04 (4.55) | 39.02 (7.28) | **0.0093** | **0.0277** |
| Linoleic | 32.46 (4.12) | 26.28 (6.74) | **<0.0001** | **<0.0001** |
| γ-linoleic | 0.48 (0.17) | 0.58 (0.18) | 0.8717 | 1.0000 |
| α-linoleic | 0.66 (0.15) | 0.62 (0.22) | 0.9533 | 1.0000 |
| Eicosadienoic | 0.19 (0.04) | 0.18 (0.02) | 0.9845 | 1.0000 |
| Dihomo-γ-linolenic | 1.71 (0.47) | 1.55 (0.35) | 0.7998 | 1.0000 |
| Arachidonic[1] | 6.98 (1.65) | 6.50 (2.05) | 0.2948 | 0.9995 |
| Eicosapentaenoic[1] | 0.58 (0.68) | 0.41 (0.16) | 0.8227 | 1.0000 |
| Adrenic | 0.25 (0.07) | 0.31 (0.07) | 0.9143 | 1.0000 |
| Docosapentaenoic n6 | 0.16 (0.05) | 0.20 (0.08) | 0.9442 | 1.0000 |
| Docosapentaenoic n3 | 0.42 (0.07) | 0.45 (0.15) | 0.9639 | 1.0000 |
| Docosahexaneoic | 2.15 (0.57) | 1.93 (0.55) | 0.7194 | 1.0000 |

[1]Non-parametric Mann-Whitney Test.
Bolded = p<0.05

**Supplemental Table 5.** Comparison of plasma fatty acids abundance (% Area) between CP subjects with and without diabetes

| **Fatty Acids** | **CP without Diabetes (n=26) Mean (St. Dev)** | **CP with Diabetes (n=21) Mean (St. Dev)** | **Unadjusted p-value** | **Adjusted p-value** |
|---|---|---|---|---|
| ***SFAs*** | 34.32 (2.85) | 35.43 (2.16) | 0.4651 | 0.8470 |
| Lauric[1] | 0.20 (0.22) | 0.36 (0.40) | **0.0417** | 0.6082 |
| Myristic[1] | 1.71 (0.51) | 1.90 (0.65) | 0.3867 | 1.0000 |
| Palmitic | 23.28 (2.64) | 23.82 (1.85) | 0.2658 | 0.9989 |
| Stearic | 7.18 (0.70) | 7.23 (0.75) | 0.9177 | 1.0000 |
| Arachidic | 0.42 (0.13) | 0.48 (0.12) | 0.8964 | 1.0000 |
| Behenic | 0.79 (0.27) | 0.84 (0.24) | 0.9182 | 1.0000 |
| Lignoceric | 0.74 (0.22) | 0.79 (0.25) | 0.9214 | 1.0000 |
| ***MUFAs*** | 27.40 (6.49) | 24.72 (3.73) | 0.0791 | 0.2190 |
| Palmitoleic[1] | 3.19 (1.98) | 2.46 (1.25) | 0.2617 | 0.9987 |
| Oleic | 22.10 (4.49) | 20.22 (2.97) | **0.0001** | **0.0022** |
| Vaccenic[1] | 1.98 (0.57) | 1.90 (0.38) | 0.9704 | 1.0000 |
| Gondoic | 0.14 (0.04) | 0.14 (0.04) | 0.9972 | 1.0000 |
| ***PUFAs***[1] | 38.28 (7.75) | 39.85 (4.82) | 0.3024 | 0.6605 |
| Linoleic | 25.45 (6.81) | 27.05 (3.78) | **0.0012** | **0.0261** |
| γ-linoleic | 0.58 (0.22) | 0.51 (0.18) | 0.8873 | 1.0000 |
| α-linoleic[1] | 0.57 (0.20) | 0.51 (0.20) | 0.7066 | 1.0000 |
| Eicosadienoic[1] | 0.17 (0.03) | 0.20 (0.04) | 0.1278 | 0.9506 |
| Dihomo-γ-linolenic | 1.47 (0.34) | 1.51 (0.38) | 0.9346 | 1.0000 |
| Arachidonic | 6.75 (1.88) | 6.66 (2.09) | 0.8600 | 1.0000 |
| Eicosapentaenoic[1] | 0.45 (0.21) | 0.41 (0.19) | 0.6676 | 1.0000 |
| Adrenic | 0.33 (0.07) | 0.30 (0.07) | 0.9523 | 1.0000 |
| Docosapentaenoic n6[1] | 0.19 (0.08) | 0.17 (0.08) | 0.4471 | 1.0000 |
| Docosapentaenoic n3 | 0.48 (0.12) | 0.44 (0.13) | 0.9446 | 1.0000 |
| Docosahexaneoic | 1.85 (0.62) | 2.01 (0.52) | 0.7389 | 1.0000 |

[1]Non-parametric Mann-Whitney Test.
Bolded = $p<0.05$

**Supplemental Table 6.** Comparison of plasma fatty acids abundance (% Area) between CP subjects with and without EPD

| Fatty Acids | CP without EPD (n=18) Mean (St. Dev) | CP with EPD (n=29) Mean (St. Dev) | Unadjusted p-value | Adjusted p-value |
|---|---|---|---|---|
| ***SFAs*** | 34.66 (2.67) | 34.91 (2.59) | 0.8713 | 0.9979 |
| Lauric[1] | 0.29 (0.39) | 0.26 (0.27) | 0.9008 | 1.0000 |
| Myristic | 1.77 (0.52) | 1.81 (0.62) | 0.9390 | 1.0000 |
| Palmitic | 23.58 (2.42) | 23.49 (2.29) | 0.8456 | 1.0000 |
| Stearic | 7.11 (0.71) | 7.26 (0.72) | 0.7683 | 1.0000 |
| Arachidic | 0.42 (0.13) | 0.47 (0.12) | 0.9215 | 1.0000 |
| Behenic | 0.76 (0.25) | 0.85 (0.25) | 0.8577 | 1.0000 |
| Lignoceric | 0.72 (0.23) | 0.79 (0.23) | 0.8980 | 1.0000 |
| ***MUFAs*** | 26.94 (5.91) | 25.75 (5.36) | 0.4496 | 0.8333 |
| Palmitoleic[1] | 2.91 (1.85) | 2.83 (1.66) | 0.9784 | 1.0000 |
| Oleic[1] | 21.90 (4.14) | 20.86 (3.82) | 0.3096 | 0.9997 |
| Vaccenic[1] | 1.91 (0.48) | 1.93 (0.50) | 0.7247 | 1.0000 |
| Gondoic[1] | 0.15 (0.04) | 0.13 (0.04) | 0.2534 | 0.9984 |
| ***PUFAs*[1]** | 38.40 (7.08) | 39.34 (6.36) | 0.7167 | 0.9773 |
| Linoleic | 25.40 (6.20) | 26.64 (5.36) | **0.0143** | 0.2716 |
| γ-linoleic | 0.52 (0.19 | 0.57 (0.21) | 0.9172 | 1.0000 |
| α-linoleic[1] | 0.62 (0.20) | 0.56 (0.19) | 0.3701 | 1.0000 |
| Eicosadienoic | 0.18 (0.04) | 0.18 (0.03) | 0.9988 | 1.0000 |
| Dihomo-γ-linolenic | 1.39 (0.31) | 1.55 (0.37) | 0.7570 | 1.0000 |
| Arachidonic | 6.89 (2.12) | 6.60 (1.88) | 0.5646 | 1.0000 |
| Eicosapentaenoic[1] | 0.46 (0.24) | 0.42 (0.18) | 0.7657 | 1.0000 |
| Adrenic | 0.32 (0.09) | 0.31 (0.06) | 0.9878 | 1.0000 |
| Docosapentaenoic n6 | 0.20 (0.10) | 0.18 (0.06) | 0.9722 | 1.0000 |
| Docosapentaenoic n3 | 0.47 (0.11) | 0.45 (0.13) | 0.9703 | 1.0000 |
| Docosahexaneoic | 1.98 (0.72) | 1.89 (0.47) | 0.8628 | 1.0000 |

[1]Non-parametric Mann-Whitney Test.
Bolded = $p<0.05$

**Supplemental Table 7.** Comparison of plasma fatty acids abundance (% Area) between CP subjects with and without Alcohol Use

| Fatty Acids | CP without Alcohol Use (n=22) Mean (St. Dev) | CP with Alcohol Use (n=21) Mean (St. Dev) | Unadjusted p-value | Adjusted p-value |
|---|---|---|---|---|
| ***SFAs*** | 34.96 (2.33) | 34.56 (2.99) | 0.8030 | 0.9924 |
| Lauric[1] | 0.23 (0.23) | 0.32 (0.41) | 0.9090 | 1.000 |
| Myristic[1] | 1.91 (0.60) | 1.67 (0.58) | 0.1773 | 0.9863 |
| Palmitic | 23.57 (2.33) | 23.45 (2.50) | 0.8125 | 1.000 |
| Stearic | 7.15 (0.83) | 7.21 (0.60) | 0.9005 | 1.000 |
| Arachidic | 0.47 (0.15) | 0.42 (0.11) | 0.9207 | 1.000 |
| Behenic | 0.86 (0.26) | 0.76 (0.26) | 0.8529 | 1.000 |
| Lignoceric | 0.78 (0.24) | 0.74 (0.24) | 0.9417 | 1.000 |
| ***MUFAs*** | 25.12 (4.03) | 27.93 (6.84) | 0.0845 | 0.2327 |
| Palmitoleic[1] | 2.46 (1.10) | 3.39 (2.18) | 0.220 | 0.9958 |
| Oleic | 23.69 (3.17) | 22.33 (4.69) | **0.0016** | **0.0346** |
| Vaccenic[1] | 1.84 (0.37) | 2.07 (0.61) | 0.3821 | 1.0000 |
| Gondoic[1] | 0.14 (0.04) | 0.14 (0.05) | 0.4714 | 1.000 |
| ***PUFAs*** | 39.92 (5.14) | 37.52 (8.12) | 0.1394 | 0.3626 |
| Linoleic | 27.33 (4.32) | 24.69 (6.69) | **<0.0001** | **<0.0001** |
| γ-linoleic | 0.53 (0.18) | 0.58 (0.22) | 0.9233 | 1.000 |
| α-linoleic[1] | 0.56 (0.17) | 0.60 (0.24) | 0.7774 | 1.000 |
| Eicosadienoic | 0.19 (0.03) | 0.17 (0.03) | 0.9741 | 1.000 |
| Dihomo-γ-linolenic | 1.58 (0.34) | 1.39 (0.35) | 0.7134 | 1.000 |
| Arachidonic | 6.42 (1.54) | 6.83 (2.41) | 0.4305 | 1.000 |
| Eicosapentaenoic[1] | 0.41 (0.20) | 0.47 (0.22) | 0.3315 | 0.9999 |
| Adrenic | 0.32 (0.08) | 0.30 (0.07) | 0.9812 | 1.000 |
| Docosapentaenoic n6[1] | 0.18 (0.07) | 0.18 (0.08) | 0.6340 | 1.000 |
| Docosapentaenoic n3 | 0.46 (0.14) | 0.47 (0.11) | 0.9753 | 1.000 |
| Docosahexaneoic | 1.96 (0.54) | 1.84 (0.65) | 0.8200 | 1.000 |

[1]Non-parametric Mann-Whitney Test.
Bolded = p<0.05

**Supplemental Table 7.** Comparison of plasma fatty acids abundance (% Area) between CP subjects with and without Smoking

| Fatty Acids | CP without Smoking (n=17) Mean (St. Dev) | CP with Smoking (n=30) Mean (St. Dev) | Unadjusted p-value | Adjusted p-value |
|---|---|---|---|---|
| ***SFAs*** | 34.44 (2.17) | 35.02 (2.82) | 0.7086 | 0.9753 |
| Lauric[1] | 0.15 (0.10) | 0.34 (0.38) | 0.1766 | 0.9861 |
| Myristic[1] | 1.80 (0.55) | 1.79 (0.61) | 0.7548 | 1.0000 |
| Palmitic | 23.16 (2.40) | 23.73 (2.28) | 0.2468 | 0.9980 |
| Stearic | 7.19 (0.85) | 7.20 (0.64) | 0.9874 | 1.0000 |
| Arachidic | 0.47 (0.12) | 0.44 (0.13) | 0.9564 | 1.0000 |
| Behenic | 0.87 (0.24) | 0.79 (0.26) | 0.8619 | 1.0000 |
| Lignoceric | 0.81 (0.22) | 0.74 (0.24) | 0.8934 | 1.0000 |
| ***MUFAs*** | 24.70 (4.49) | 27.05 (5.97) | 0.1342 | 0.3510 |
| Palmitoleic[1] | 2.01 (0.65) | 1.87 (0.53) | 0.6256 | 1.0000 |
| Oleic[1] | 20.54 (3.27) | 21.66 (4.30) | 0.5798 | 1.0000 |
| Vaccenic[1] | 1.77 (0.45) | 2.04 (0.49) | 0.0555 | 0.7155 |
| Gondoic | 0.14 (0.04) | 0.14 (0.05) | 0.9941 | 1.0000 |
| ***PUFAs*** | 40.85 (5.29) | 37.92 (7.08) | 0.0624 | 0.1758 |
| Linoleic | 27.33 (5.33) | 25.51 (5.83) | **0.0003** | **0.0066** |
| γ-linoleic | 0.56 (0.21) | 0.54 (0.20) | 0.9685 | 1.0000 |
| α-linoleic | 0.54 (0.16) | 0.61 (0.22) | 0.8857 | 1.0000 |
| Eicosadienoic | 0.18 (0.03) | 0.19 (0.38) | 0.9781 | 1.0000 |
| Dihomo-γ-linolenic | 1.45 (0.31) | 1.50 (0.38) | 0.9198 | 1.0000 |
| Arachidonic | 7.31 (1.7) | 6.36 (1.99) | 0.0558 | 0.7172 |
| Eicosapentaenoic | 0.48 (0.23) | 0.41 (0.19) | 0.8815 | 1.0000 |
| Adrenic | 0.32 (0.08) | 0.31 (0.07) | 0.9841 | 1.0000 |
| Docosapentaenoic n6[1] | 0.18 (0.09) | 0.19 (0.08) | 0.6806 | 1.0000 |
| Docosapentaenoic n3 | 0.51 (0.11) | 0.44 (0.13) | 0.8849 | 1.0000 |
| Docosahexaneoic | 2.01 (0.65) | 1.87 (0.53) | 0.7822 | 1.0000 |

[1]Non-parametric Mann-Whitney Test.
Bolded = p<0.05
